# Directionality and community-level selection

**DOI:** 10.1101/484576

**Authors:** Guy Bunin

## Abstract

Many ecological community dynamics display some degree of directionality, known as succession patterns. But complex interaction networks frequently tend to non-directional dynamics such as chaos, unless additional structures or mechanisms impose some form of, often fragile or shot-lived, directionality. We exhibit here a novel property of emergent long-lasting directionality in competitive communities, which relies on very minimal assumptions.

We model communities where each species has a few strong competitive interactions, and many weak ones. We find that, at high enough diversity, the dynamics become directional, meaning that the community state can be characterized by a function that increases in time, which we call “maturity”. In the presence of noise, the community composition changes toward increasingly stable and productive states. This scenario occupies a middle ground between deterministic succession and purely random species associations: there are many overlapping stable states, with stochastic transitions, that are nevertheless biased in a particular direction. When a spatial dimension is added in the form of a meta-community, higher-maturity community states are able to expand in space, replacing others by (exact or approximate) copies of themselves. This leads to community-level selection, with the same maturity function acting as fitness. Classic concepts from evolutionary dynamics provide a powerful analogy to understand this strictly ecological, community-level phenomenon of emergent directionality.

---

A wide range of spatio-temporal patterns in ecological communities exhibit some degree of directionality and predictability, and are studied collectively under the name of succession patterns [39].

Ecological succession was originally pictured as a predictable sequence of community compositions, akin to an organism’s developmental stages [14]. But this simple and enticing picture was rapidly shown to present two major discrepancies with natural patterns. First, communities rarely display only a few discrete well-separated states. Instead, species are found in numerous overlapping combinations, leading many ecologists to reject the very concept of stable species association [24].

Second, changes in community composition are not necessarily fully deterministic or sequential. Some communities, from bacteria to forests, exhibit stochastic [25] and chaotic [8] changes in dominance. Complex and nondirectional dynamics, from cycles to chaos, are found in many-species dynamical models in ecology [7, 26], evolution [17] and game theory [16, 23].

Yet, a single ingredient is needed to recover a picture of ecological succession that is directional, yet allows many species associations and stochastic trajectories. This scenario can occur when species interact strongly with some competitors, and weakly (if at all) with others (the “many weak, few strong” scenario, e.g., [52, 55]). Such mutually exclusive interactions have been observed in lab experiments due to multiple mechanisms [1, 37, 56]; while their effect may be harder to observe in nature, they show up in the form of spatial heterogeneities [15, 47] and priority effects [22]. They play a major role in shaping these communities and their response to invasion, a picture known as “the ghost of competition present” [38].

We show theoretically that this hypothesis provides a robust mechanism for the emergence of directional ecological dynamics. Thus, it brings unexpected insight into the (seemingly unrelated) question of ecological succession.

In this setting, a community can be driven into many alternative states which represent different, and often overlapping, combinations of the same species [20, 21]. We find that each of these associations can be characterized by a single quantity, which we call its maturity. As one increases both the species diversity and the intensity of the strong competitive links, jumps become more and more biased toward higher maturity states.

This mechanism creates an intermediate situation between sequential community stages and independent species dynamics. Here, succession is not predetermined or imposed, but an emergent many-species behavior, with many paths leading, in a stochastic fashion, toward increasingly mature communities.

This mechanism allows for dynamics over time-scales that are much longer than generation or colonization time, since the system remains in more mature states for longer times (a form of stability). This is in contrast to a picture of succession driven solely by how long a species takes to grow.

We find that a community’s maturity is strongly associated with functional properties such as total productivity, which we predict to increase in long-term dynamics. This ties our results to the functional approach of ecological succession championed by Margalef [35] and Odum [39]: while changes in composition do not follow a simple and fully deterministic sequence, changes in functioning tend to be predictable.

This scenario also paints a compelling picture of spatial heterogeneity and dynamics. The existence of many alternative but overlapping subsets of the same species roster provides a biotic explanation for spatial diversity and patchiness in macroscopically-homogeneous ecosystems such as grasslands [31].

Most strikingly, we show in this spatial context that the maturity of a community state also predicts its tendency to spread to adjacent patches. Here, concepts from evolutionary dynamics provide a powerful analogy. Even though all the mechanisms involved are strictly ecological (specifically, species’ interactions and dispersal), states that expand efficiently in space can take over the landscape, in analogy with the way that high-fitness genotypes (or phenotypes) are selected for in evolutionary population dynamics. Thus, directionality may be enhanced by the spatial dimension through a community-level selection process. This surprising mapping is helpful in understanding the middle ground we suggest for succession: in evolution, the specific progression by which mutations accumulate may be stochastic, but overall the fitness is expected to increase. Here similarly, the species associations may change stochastically but maturity will nonetheless increase.

The paper is structured as follows: Sec. I describes the model for a single community of interacting species without spatial structure. We introduce the essential ingredient of the model, see Fig. 1(A).

**Figure 1.**
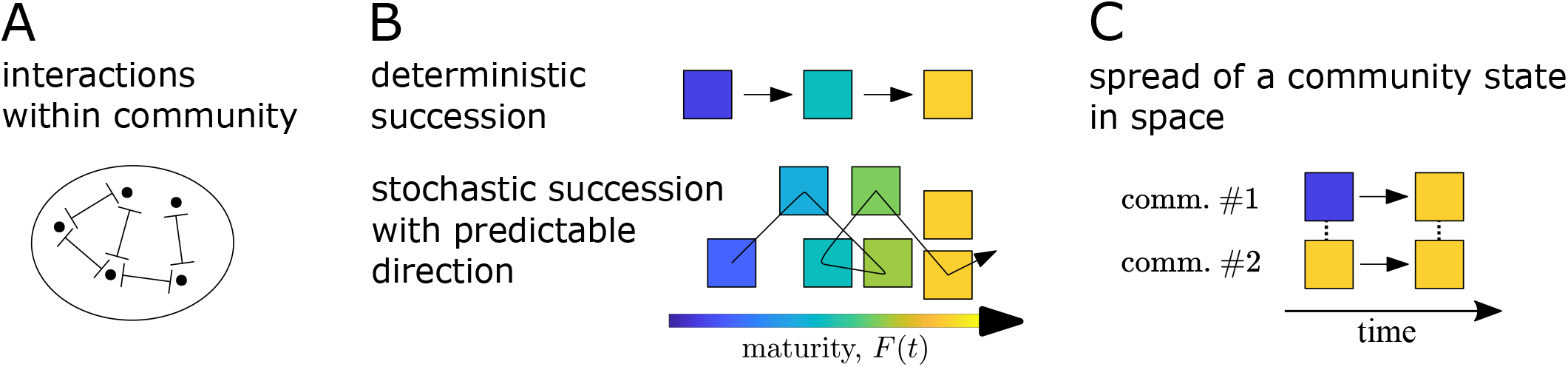
(A) Illustration of a community in which some pairs of species are mutually exclusive (marked by 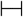 symbols). Other interactions may also be present. (B) A known function, the *maturity F*, of the community imposes directionality in the dynamics of the system. The ecological community exhibits multiple alternative states (squares, signifying different possible community compositions). Changes in time between states are not fully deterministic (top) but stochastic, yet with a tendency to transition towards states with higher values of *F*. (C) When multiple local communities are coupled by migration (dashed lines), certain states are found to expand across the different communities. Here, the state of community #1 changes at some time, to match that of community #2. States with higher values of maturity *F* tend to expand more efficiently.

Sec. II,III,IV, discuss how directionality emerges at sufficiently high species diversity, and describe the function that increases in time, see Fig. 1(B). The same function increases also in the presence of noise, as the system moves between multiple alternative states, exploring ever more stable, longer lasting states. The ecological interpretation of this function and its relation to production are discussed.

In Sec. V, insight into these phenomena is obtained from a simple instance of the general model, with just 3 species. Spatial structure is then added to the model, realized as a meta-community, in which multiple communities are coupled by migration. The simple model demonstrates some of the phenomena that appear in high-diversity cases: that states may expand in space and that the same function that increases in a local community is also associated with better ability to replicate, and thus acts as a fitness function, see Fig. 1(C). This contributes to directionality by selection of higher values of the function, in analogy with the increase of fitness in population dynamics. In Sec. VI, these results are then shown to hold and generalize to the full, high-diversity spatial model. Sec. VII discusses the generality and limitations of the model, followed by a discussion in Sec. VIII.

## I. MODEL DEFINITION

Here a single, well-mixed community is discussed, with spatial structure left for later sections. Motivated by the “few strong, many weak” scenario (e.g., Vázquez et al. [52], Wootton and Stouffer [55]), we describe an ecological community, with predominantly competitive interactions, and where every species is, on average, *mutually exclusive* with a few other species. By this we mean that if these two species are mixed, a stable equilibrium will contain one or the other species, but not both. Other interactions are weak or non-existent. Many model variants will give similar results, for concreteness we focus on the usual Lotka-Volterra model for the relative abundances (yields) of *S* species, *N_i_*_=1..*S*_,

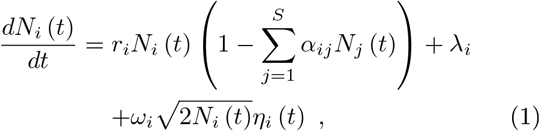

with *α_ii_* = 1. The *λ_i_* are time-independent constants that account for migration (difference between immigration and emigration). In the demographic noise term, 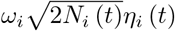, *η_i_* (*t*) is random time-varying function, and here we use a Gaussian process with white noise. The 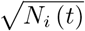 factor is chosen to capture the scaling of demographic noise with population size [30]. The terms *ω_i_* are time-independent constants, setting the overall strength of the noise, which relates to the absolute number of individuals of each species: larger populations experience smaller demographic noise on the relative abundance. The *r_i_* terms are time-independent growth rates, and *α_ij_* terms are time-independent interaction coefficients. All definitions, parameter values and simulation details for the entire paper are described in Materials and Methods, Section IX.

In Eq. (1), mutual exclusion translates to a pair *i, j* for which both *α_ij_, α_ji_* > 1. In the following, the weak interactions are set to zero so the matrix *α* is sparse. Motivated by niche theory where species interact when their niches overlap, it is assumed that *α_ij_, α_ji_* are either both non-zero or both zero. *α_ij_, α_ji_* are not assumed to be equal or similar in their values. Each species interacts with *C* other species. Additional model parameters are given in the following sections.

In order to disentangle different effects, other processes are excluded, such as evolution (leading to changes in interactions, speciation, etc.), additional community replication [50] (e.g., by host replication [42]) or external selection of entire communities [53]. The focus is on the consequences of this minimal setting, rather than on the possible biological contexts that might lead to this behavior. In this way it uncovers the crucial ingredients that allow this behavior to be robust, see Sec. VII, such as replacing the zero-values interactions with sufficiently weak interactions.

## II. DIRECTIONALITY AND STABILITY IN A MULTI-STABLE SETTING

In this section we show how directionality emerges in many-species dynamics, in the presence of strong competition.

Other models have imposed directionality, e.g. certain resource competition models [33, 34], or models with specific constraints as discussed below. Here we show that, even if it is not incorporated in the model by construction, directionality can emerge dynamically provided that even a small fraction of interactions are sufficiently competitive.

We first consider the noiseless case, i.e. setting all *ω_i_* = 0. When the matrix *A* with *A_ij_* ≡ *r_i_α_ij_* is symmetric it is well known (e.g., [40]) that the function

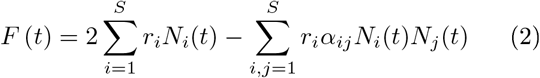

is non-decreasing during the dynamics of Eq. (1) without noise. Such a function is known as a Lyapunov function. Yet, if *A* is not symmetric^1^ *F* may generally decrease during the dynamics. Let’s look at what happens in this more general case.

Fig. 2(A), shows runs with different values of mean (*α_ij_*), the mean of all non-zero interactions, keeping all other parameters constant (parameters are given in the Supporting Information, Appendix IX). When mean (*α_ij_*) is small enough such that most interactions satisfy *α_ij_* < 1, *F* (*t*) first increases and then fluctuates indefinitely, indicating that the abundances fluctuate, and the dynamics do not reach a fixed point^2^. Increasing mean (*α_ij_*), we see that systems are more likely to converge to fixed points, after some oscillations of the function *F* (*t*). Increasing the mean further, so that for a large fraction of the non-zero interactions *α_ij_* > 1, the function *F* (*t*) *increases throughout the entire dynamics*. In other words, *F* (*t*) behaves as if it were a Lyapunov function^3^. It always converges to a fixed point. Indeed, the fact that *F* increases precludes persistent abundance fluctuations, as those shown in Fig. 2(A) for low values of mean(*α_ij_*).

**Figure 2.**
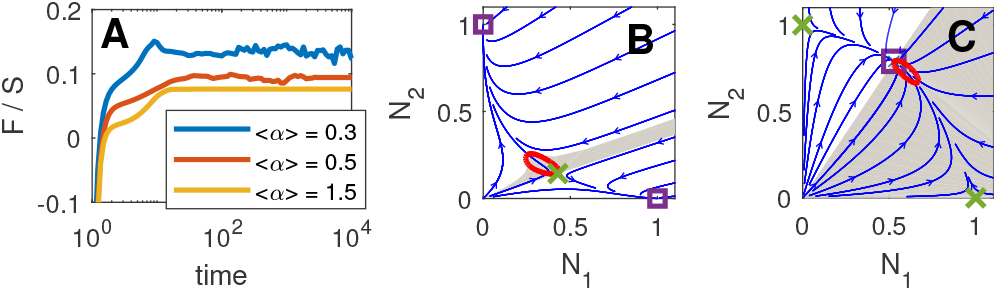
Directionality can emerge, even if not imposed by construction. (A) The function *F* (*t*) in a system with many species, for three different runs. In each run the mean value of the non-zero *α_ij_* is different, with other parameters kept constant. As the mean is increased, the behavior changes from persistent fluctuations in *F* (*t*) and dynamics that do not reach a fixed point, to dynamics in which *F* (*t*) only increases until equilibrium. Comparison between mutual-exclusion (B) and coexistence (C) for *S* = 2, gives an idea for the reasons for the growth of *F*. *F* (*t*) decreases when entering the areas delineated in red, which happens for dynamics initialized any - where in the gray regions. Squares and crosses mark stable and unstable fixed-points, respectively. In (A), each species interact with *C* = 20 other species, and *S* = 400.

As *F* (*t*) increases in time, both here and with the introduction of noise and spatial structure below, we will call it the *maturity*. This reflects the ability of this function to distinguish between earlier and later states: if two states appear in the history of a community, a directional function allows us to determine which appear later (the state with higher value of *F*), thus acting as a measure of how mature a system is. The ecological interpretation of the function *F*, as well as a more suggestive expression than Eq. (2), is discussed in Section IV.

## III. NOISE AND TRANSITIONS TO LONGER-LASTING STATES

In the presence of noise the system may switch between otherwise stable states. The term ‘state’ refers to a stable equilibrium together with its basin of attraction under noiseless dynamics. Fig. 3 shows results for noisy dynamics. Here and in Sec. IV,VI, the species pool has *S* = 24 species, each interacts with *C* = 3 other species, and the non-zero *α_ij_* are sampled from a Gaussian distribution with mean (*α_ij_*) = 1.667 and std (*α_ij_*) = 0.75. With these parameters, all species pairs are either competitive (*α_ij_* > 0) or do not interact. The growth rates *r_i_* are sampled uniformly in (0.5, 2). Other parameters are given in the Supporting Information, Appendix IX.

**Figure 3.**
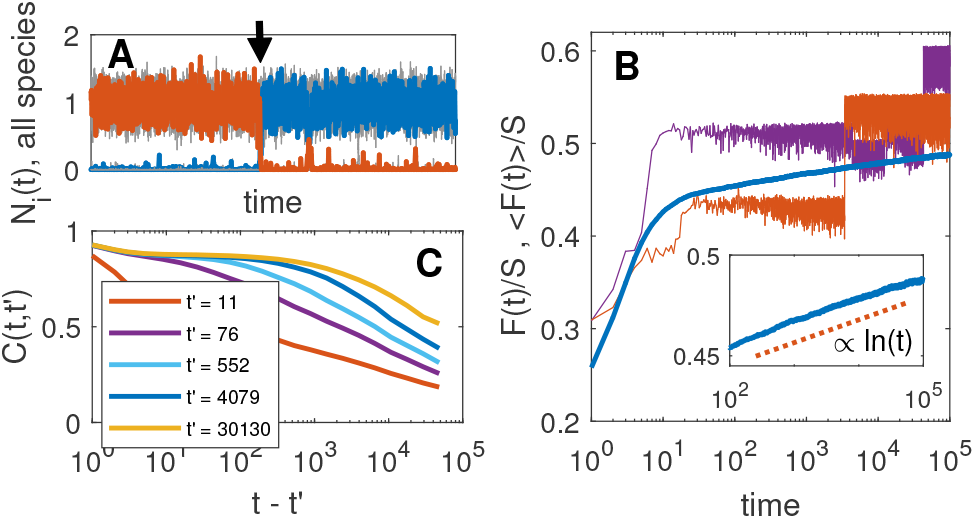
Transitions between states in noisy dynamics. (A) Each species is either found at very low abundance maintained only by migration, or is established. The system may undergo noise-driven changes of state, as in the simple one marked by an arrow, where one established species is replaced by another. (B) The function *F* (*t*) in two example runs (thin lines), and the mean over many runs (thick line). Inset: close up on the mean *F* (*t*) at later times (*t* > 100), showing its continued growth. (C) The correlation function between two times *t*′ and *t* remains high for longer times as *t*′ grows, showing that the dynamics switch states less frequently as more stable states are explored. Here and in Figs. 4,6, each species interacts with *C* = 3 other species, and *S* = 24.

Fig. 3(A) shows an example of a transition in the composition of species, where one species is able to invade at the expense of another which is removed from the community. As expected, the maturity *F* (*t*) fluctuates due to the noise; remarkably it tends to *increase* in time. It is shown in Fig. 3(B) for two example trajectories. At long times, *t* ≳ 100, *F* (*t*) shows periods of fluctuations within a single state, punctuated by abrupt transitions in which *F* (*t*) more often increases than decreases. The mean of *F* (*t*) over many runs increases at long times roughly as ⟨*F* (*t*)⟩ ~ ln *t*, see Fig. 3(B). The angular brackets ⟨..⟩ denote average over samples of the interactions, growth rates, initial conditions and noise. This growth at long times is a non-trivial behavior, which is not found in many other models or parameter regimes, such as when noise is added to the chaotic regime discussed above, causing ⟨*F* (*t*)⟩ to quickly reach a plateau (Supporting Information, Appendix A).

With time, the dynamics spend longer periods of time in each state, as can be appreciated from the runs shown in Fig. 3(B) (note the logarithmic scale in the time axis). To quantify whether the state has changed, and if so by how much, we look at the correlation function *C* (*t, t*′), measuring the similarity (correlation coefficient) between the abundances *N_i_* (*t*′) and *N_i_* (*t*), with *t*′ < *t*. Fig. 3(C) shows that at later times *t*′, the correlation remains high value for larger time differences *t* − *t*′, reflecting a *growth in stability*, in the sense that the system spends ever longer times in any given state or in states similar to it^4^. Thus, states with larger *F* are also more stable.

The transitions between states appear as jumps in Fig. 3(A,B), demonstrating that the system has multiple equilibria. There are various ways in which a highly-diverse community may admit multiple-equilibria [10, 13, 16, 20, 21], resulting in different organizational and dynamical properties. The structure of equilibria in communities whose interactions are predominately mutually-exclusive has been studied in [20, 21]. Multiple equilibria are expected to appear in this setting, and for large systems very many of them – exponentially many in the number of species *S* [21], as we also find in the present model, see Supporting Information Appendix B.

The bias towards higher *F* in the dynamics provides a simple mechanism for the growth in stability, as transitions are biased towards states with higher *F*, so that states with large *F* have less such states to switch to.

## IV. THE MATURITY *F* (*t*) AND GROWTH OF PRODUCTION

Over longer times, the changes in *F* (*t*) slow down, and *F* (*t*) is approximately equal to an alternative and more suggestive form [32].

When the dynamics fluctuate around a given equilibrium, and after averaging out the rapid fluctuations due to the noise, changes in the total biomass will vanish, 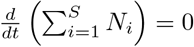 Eq. (1) reads

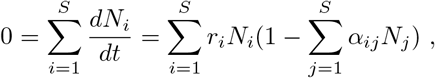

where migration is assumed to be negligibly small. Rearranging and using Eq. (2) we obtain 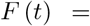 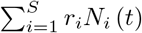. If instead the system does change over time but slowly, so that 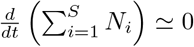 we find only an approximate equality,

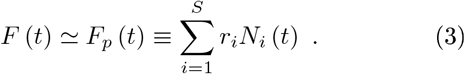

The expression for *F_p_* (*t*) in Eq. (3) is simply the average over bare growth rates *r_i_*, weighted by the abundances *N_i_* (*t*). It is thus a form of production [32] – it would be equal to the total biomass production if the growth rates corresponded to the per-capita biomass production. As shown in Fig. 4(A), *F_p_* (*t*) and the maturity *F* (*t*) both fluctuate around a common average, which moves upon transitions between states. The averages over many realizations of the model, ⟨*F_p_* (*t*)⟩, ⟨*F* (*t*)⟩, match closely. *F* (*t*) can be seen to fluctuate less than *F_p_* (*t*) in Fig. 4(A), since *F* (*t*) fluctuates around a local maximum, while *F_p_* (*t*) is a simple linear function of the abundances, which does not admit a local maxima at the equilibria. *F_p_* (*t*) does not generally increase when equilibria are approached within the basin of a single equilibrium, so while stability of states and the transitions between them are closely tied to the increase in the maturity *F* (*t*), the increase in the production *F_p_* (*t*) is in a sense a byproduct of the behavior of *F* (*t*).

**Figure 4.**
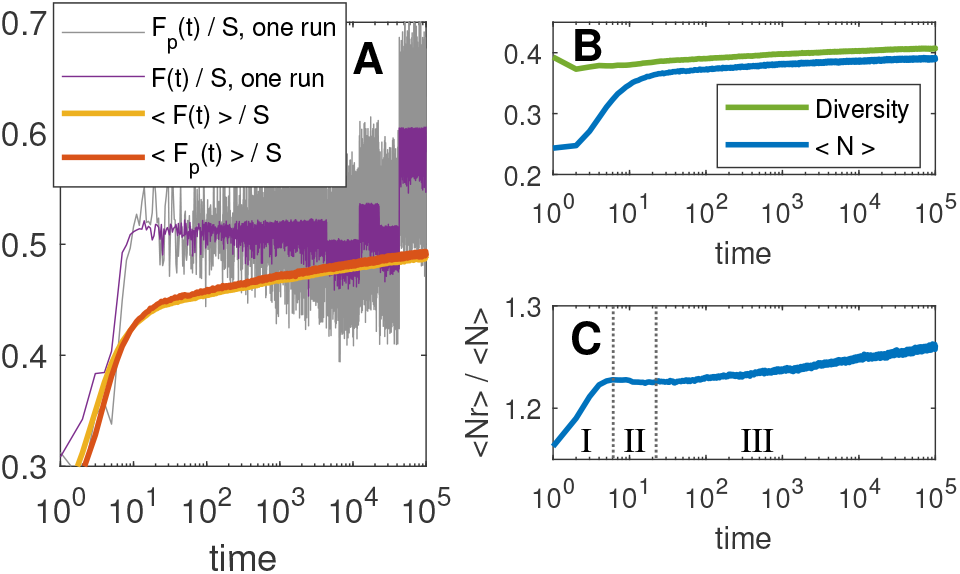
Growth of production. (A) Comparison of maturity *F* (*t*) and production *F_p_* (*t*). Over one run (thin lines), they fluctuate around the same values, with *F_p_* (*t*) showing larger fluctuations. The averages over many runs match well. (B) Over long times, the diversity grows (the fraction of established species, *N_i_* > 0.25, see Fig. 3(A)), as does the average abundance. Apparent early decline in diversity comes before first state stabilizes (see also Fig. 6(D)), at which times the diversity criteria for isn’t relevant. (C) Mean per capita growth rate, showing three periods: I. colonization (increase of function); II. maturation within state (saturation and some decrease); III. jumps between states (increase, again). growth rate, showing three periods: I. colonization (increase of function); II. maturation within state (saturation and some decrease); III. jumps between states (increase, again).

The function *F_p_* (*t*) can be split into the product of the change in the total abundance Σ_*i*_*N_i_* (*t*) and the weighted average of the growth rates, Σ_*i*_*N_i_r_i_*/Σ_*i*_ *N_i_*, which is akin to productivity [39] (biomass production per unit biomass). In simulations of the model, the later is found to have a characteristic shape, see Fig. 4(C): at very short times it grows rapidly, as species with large *r_i_* increase in abundance in this colonization period. It then levels off and decreases a little. This saturation and decrease in productivity is known to occur during community maturation [39], as fast growing species (high *r_i_*) are replaced with species that are able to coexist. At later times (*t* ≳ 100) the community transitions between different possible states. At this stage the diversity of the community, the average growth rate and the total abundance Σ_*i*_*N_i_* all gradually increase, see Fig. 4(B). These can be related the growth of *F_p_* (*t*), with the growth of Σ_*i*_*N_i_r_i_* entailing that of averages of *N_i_* and *r_i_*. It might be more model dependent.

## V. STABILITY, DIRECTIONALITY AND SPATIAL EXPANSION–A THREE-SPECIES TOY EXAMPLE

To get some intuition on the growth in production *F_p_* and stability in the presence of noise, we study a simple instance of the model. We then use it to introduce spatial structure, before adding spatial structure to the full, high-diversity setting in the next section. The simple model features three species. Species 1 has mutual exclusive relations with 2 and 3, see Fig. 5, and species 2,3 have no direct interaction (they could interact sufficiently weakly without changing the results). In this situation there are two equilibria of the noiseless dynamics: one with species 1 present and 2,3 suppressed due to the mutual exclusive relations; and another with 2,3 present and 1 suppressed. They will be denoted by *s*_1_ and *s*_23_ respectively. We take all *r_i_* = 1, returning to that later.

**Figure 5.**
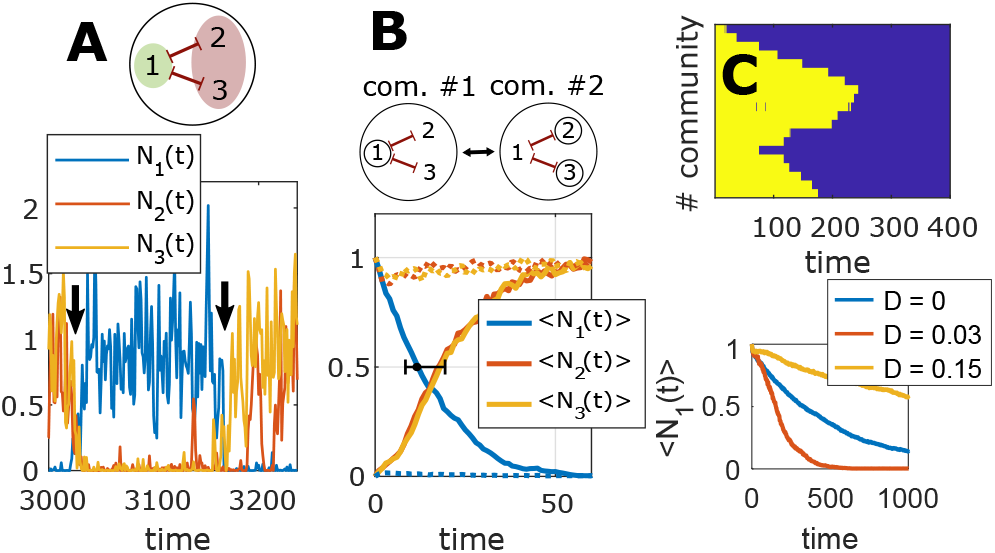
(A) A simple community, with three species (1,2,3) and two competitive-exclusion relations (marked by 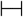 symbols), with *α_ij_* = *α_ji_* = 1.5. This has two equilibria: one (*s*_1_) with only species 1 present and 2,3 suppressed, and the other (*s*_23_) with species 2,3 present, and 1 suppressed (colored ellipses). The system jumps more easily from the state *s*_1_ to the state *s*_23_. Both transitions are shown in an example run (marked by arrows). (B) When communities are coupled by migration, states can expand to neighboring communities. Here, two communities as in A are coupled, one initialized in state *s*_1_ and the other in *s*_23_ (circled species). The state *s*_23_ rapidly takes over both communities. Shown are time averages over repeated runs, for community initialized in *s*_1_ (solid lines) and in *s*_23_ (dotted lines). The horizontal errorbar shows the standard deviation in the time when *N*_1_ crosses below 0.5. Migration rate is *D* = 0.03. (C) A chain of *M* = 20 coupled communities, all initialized in state *s*_1_ (in yellow, defined by *N*_1_ (*t*) > 0.25). The state changes to *s*_23_ in two places, and expands to all the communities (here *D* = 0.03). Bottom: ⟨*N*_1_⟩ (*t*) is shown for *M* = 20 and different *D*. The transition to *s*_23_ may be faster (*D* = 0.03 as in top panel) or slower (*D* = 0.15) than for a single community (*D* = 0). In all panels *ω_i_* = 0.173 for all species.

With noise, the system fluctuates around one of the two equilibria, with occasional transitions between them, see Fig. 5(A). The maturity-values of the two equilibria can be readily calculated from Eq. (2) or Eq. (3), giving *F*_1_ = 1 and *F*_23_ = 2. Therefore we expect the system to jump more easily from *s*_1_ to *s*_23_ than in the opposite direction, and so spend more time in *s*_23_. Indeed, with 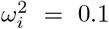 the system is at *s*_1_ only about 7% of the time (measured by the fraction of time that *N*_1_ (*t*) > *N*_2_ (*t*), *N*_3_ (*t*)).

To understand the asymmetry in the transition rates, consider the transition processes. The system switches from *s*_1_ when the abundance of *N*_1_ is low enough, by some random fluctuation driven by the noise, so that either *N*_2_ or *N*_3_ grow from their low values, see the second transition in Fig. 5(A). Transitioning in the other direction requires both *N*_2_ and *N*_3_ to be unusually low simultaneously before *N*_1_ can increase, see the first transition in Fig. 5(A). The latter transition is therefore less likely. This is a mechanism for growth in Σ_*i*_ *N_i_*, representing higher total abundance or diversity: transitioning from high- to low-diversity requires more species to be unusually low simultaneously, than vice-versa.

If species have different growth rates, a preference for higher *r*-values comes from two mechanisms. First, species with higher *r* are more stable, with smaller noise-induced fluctuations. Secondly, upon transitioning, species with higher *r*-values grow faster, making transitions to states in which they are present more likely.

We now introduce spatial structure. Space is modeled within the meta-community framework, in which *M* communities are coupled by migration, and each community is well-mixed and subject to internal dynamics as in Eq. (1). Let 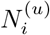 be the abundance of the *i*-th species in the *u*-th community. The equations read

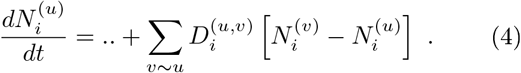

where the “..” refers to the terms in the RHS of Eq. (1), applied to 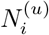. In particular the small external migration term *λ_i_* is kept in all communities, meaning that they are also weakly coupled to an external pool of species, so that all species can try to invade. Interactions *α_ij_* and growth rates *r_i_* are taken to be the same in all *M* communities. In the following we take all migration rates to be identical, 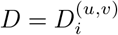.

In Fig. 5(B), two communities are coupled by migration, with the interactions in each community as discussed above (Fig. 5(A)). The abundances are initialized with one community in equilibrium *s*_23_ and the other in *s*_1_. Without noise and at sufficiently small migration rate *D*, this assignment of abundances is stable: each community is stable against the invasions via migration from the other community. However with noise, the state *s*_23_ is able to expand and take over both communities, see Fig. 5(B), which shows ⟨*N*_1,2,3_ (*t*)⟩ in the community initialized at *s*_1_ (with *N*_1_ = 1, *N*_2_ = *N*_3_ = 0 at *t* = 0), quickly transitioning to state *s*_23_. The reason for the asymmetry in the transition is again that the takeover requires only one species to decrease in *s*_1_, compared to two in the community with *s*_2_, and migration from a neighboring community catalyzes the process.

Coupling more communities allows for spreading of a state throughout the meta-community. In Fig. 5(C), 20 communities are coupled in a linear chain, each to its two nearest neighbors. The dynamics in time and space are visualized in two dimensions, with the vertical axis showing the patches as ordered along the chain, and the horizontal axis corresponding to time. The system is initialized with all communities in state *s*_1_. We see that state *s*_23_ eventually takes over. In the top example the switch first happens in two communities, and then spreads to the rest of the system via migration. This change happens faster in the meta-community than for a single community, see bottom panel.

Migration introduces two competing effects. On the one hand, migration acting between communities in the same state can reduce the chances of a transition, by suppressing changes to the community’s state. Indeed, in the limit of strong migration (high *D*) the abundances are equalized between all communities, and the metacommunity behaves as a single well-mixed community, with lower effective noise (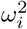 reduced by a factor of 1*/M*) due to the larger absolute populations of each species. On the other hand, with migration the transition can happen in only one community, and then expand to the entire meta-community. This increases the chances of transition per unit time at weaker migration levels. Due to the competition between the two effects, the mean time until the transition is a non-monotonic function of the migration rate *D*, see bottom panel of Fig. 5(C). Similar non-monotonic dependence was found in [29].

We showed that the state in one community is able to influence the state of communities coupled to it, such that they adopt its state. This process can be thought of as an “infection” or, when the state is a perfect copy, a “replication” process. It allows for selection at the level of the meta-community: states that are better able to replicate will be found in more communities, and in this way selected for in the population of communities. In an analogy with selection in Darwinian evolution, each patch corresponds to an individual organism in a fixed size population (the fixed number of communities). Spontaneous, noise-driven changes of a community’s state are analogous to mutations. And as states with higher *F*-values are also better replicators, the maturity *F* corresponds to *fitness*. We now turn to discuss this in the full, high-diversity setting.

## VI. META-COMMUNITIES AT HIGH DIVERSITY: COMMUNITY-LEVEL SELECTION

This section looks at meta-communities in the full, high-diversity setting. Compared to the toy model of Sec. V, here the dynamics explore a more complex space, with many equilibria that overlap to varying degrees in their species composition. It is therefore far from obvious that intuitions from a two-state model will carry through to this case. In particular, while a community in Sec. V can only jump between two states, here a state might expand in an “imperfect” way, inducing similar but not identical states in neighboring communities. We find that, as in the simple model discussed in in Sec. V, states with higher *F*-values are also better replicators, and so spatial expansion promotes the growth of the maturity *F* via a selection process.

In Fig. 6 the communities are arranged in a linear chain, as in Fig. 5(C). Fig. 6(A) shows one run. For each time and community, the value of *F* is plotted, at the equilibrium associated with each community state (found by running zero-noise dynamics of the community in isolation). A number of distinct time periods can be seen. First, at *t* ≲ 10, the communities relax into different states (see Fig 6(D)). The transitions between states are less frequent and states in different communities become more similar, and by time *t* = 300, all communities assume the same state. The cross-correlation between different communities is presented in the Supporting Information Appendix B (Fig. E). Then there is no change until *t* = 1380 at which time all communities transition to a new state with higher *F*. The change occurs within a short time frame of less than 100 time units, and so looks like a jump in *F* in Fig. 6(B). A close - up on this time frame, Fig. 6(C), shows a front of the new state which sweeps through the chain of communities, until taking over the entire system. In the analogy with population dynamics, a new state (analogous to a mutation) has spread and fixed in the population. This expansion forms copies of exactly the same state, analogous to a replication process. This is the most common mode of change at late times, see also additional runs at higher diversity (*S* = 100) in the Supporting Information Appendix B.

**Figure 6.**
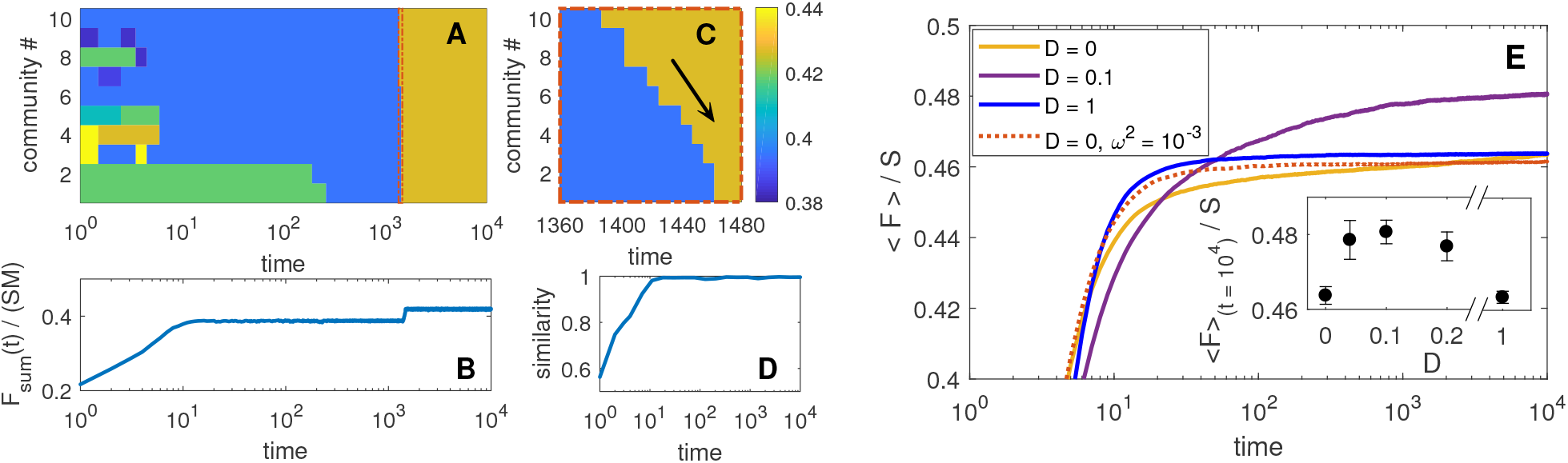
Meta-community dynamics. (A) An example run, showing the maturity *F* of the equilibrium corresponding to the state of each community (defined as *F* of the equilibria reached by zero-noise dynamics of the patch in isolation). (B) Average of *F* over all the communities (of the instantaneous states, that are not necessarily equilibria). (C) A close-up on a change in population state (narrow red rectangle in (A)). A new community state, first appearing in the top-most community, spreads until taking over the entire meta-community. The arrow shows the direction of the spread. (D) The similarity of the noisy state to the zero-noise equilibria, showing the initial dynamics towards an equilibrium, up to time *t*~10. (E) The mean ⟨*F* (*t*)⟩ over all communities and many runs, for different migration rates *D*. Intermediate values (*D* = 0.1) allow ⟨*F* (*t*)⟩ to grow well above the value reached for a single community (*D* = 0). High migration rates (*D* = 1) suppress the growth of *F* at late times, and the system behaves as if it is well-mixed with smaller noise (dotted line). Inset: ⟨*F*⟩ at time *t* = 10^4^, which depends non-monotonically on *D*. In all panels there are *M* = 10 communities, all *ω_i_* = 0.1, and other parameters as in Fig. 3. In (A-D), *D* = 0.1.

Spatial expansion leads to selection of states with higher *F*, as shown in Fig. 6(A,B,C). This allows the average ⟨*F* (*t*)⟩ to reach higher values than those attained for a single community (*D* = 0) at late times, see Fig. 6(E). The rate of any given transition between two states is expected to depend non-monotonically on the migration rate *D*, as discussed for the 3-species model in Sec. V. This translates here to a non-monotonic dependence of ⟨*F* (*t*)⟩ on *D*, see the inset in Fig. 6(E).

## VII. GENERALITY AND LIMITATIONS

The phenomena described above persists under many variations on the model details, and so might be found in more biologically-realistic models, and be relevant to natural or laboratory settings. Elements which do not change the qualitative behavior include (for details and simulation results see Supporting Information, Appendix A): some fraction of pairs *ij* where only one of the *α_ij_* is non-zero; interactions that are not of a Lotka-Volterra form (saturating interactions); niche structure, in which the pairs with non-zero interactions are chosen according to the distance in niche space; and additional interactions between all pairs, that are less competitive (not mutually exclusive). All the phenomena described do not require very high diversity: the examples in Sec. III–VI used *S* = 24 species, and are already present with even fewer species.

At the same time, once certain parameters are changed beyond some limit, the system dramatically changes its qualitative behavior. This includes changes which might at first sight seem innocuous or unrelated to the phenomena of interest. For example, directionality or growth of stability will be lost if the mean interaction strength is lower than some value (see Sec. II), or if the noise is higher than some value (Supporting Information, Appendix A).

High-diversity communities exhibit distinct qualitative regimes, known as phases, controlled by a few driving parameters [5, 13, 16, 19, 27, 49]. Identifying these sometimes counter-intuitive parameters is key to understanding the potential behaviors of a system and the transitions between them. In these terms, the present work characterizes a distinct phase.

## VIII. DISCUSSION

### Predictions

In this work, we have looked at the dynamics of ecological communities, driven by species interactions and migration. The scenario studied here is characterized, and can be identified in an ecosystem by a combination of phenomena, which impact multiple aspects of the community dynamics. They are summarized in the following list.

- The existence of mutually-exclusive pairs of species, and weaker interactions (if any) between other pairs. These can be identified by pair competition experiments.
- There are very many equilibria, and at the same time they represent only a small fraction of all possible species associations. To illustrate this, we estimate (Supporting Information, Appendix B) that for *S* = 100, with interactions sampled as in Fig. 3,4 and 6, there will be hundreds of millions of possible equilibria. This large number, does not mean that any collection of species may stably coexist, as equilibria represent only a small fraction of about 2^100^ ≃ 10^30^ possible species’ subsets.
- Relatively sharp spatial transitions between community states, even in the absence of any change in abiotic conditions, which can be maintained for some period of time. This is possible as nearby patches may remain in different uninvadable equilibria. For example, in Fig. 6(A) different patches remain in two different states during time 6 ≲ *t* ≲ 200.
- Changes in species’ composition via population sweeps, especially at late times, as shown in Fig. 6(C).
- The diversity tends to grow with time, as does the maturity function *F* (*t*). This function is interpreted as the bare productivity weighted by the yields, see Eq. (3), and can thus be calculated if bare growth rates and abundances are measured. Meta-communities may behave very differently from well-mixed communities. Under some conditions, *F* may reach significantly larger values.
- Changes in the community states slow down significantly as time passes, with the system moving ever more slowly between community states. This can be seen both through a slower changes in the maturity and diversity (see previous point), and through the correlation between the abundances at different times, as shown in Fig. 3(C).

### Emergent processes of succession

A single function (*F*) of the community species’ abundances governs two processes: first, this function increases during the dynamics of a single community, and so ensures directionality in its succession pattern. It was therefore called the maturity. Secondly, this function is correlated with the ability of a community state to expand to other communities. In turn, this expansion process enhances the directionality in meta-communities, by selection of states that efficiently expand in the population of communities, in analogy with the increase in fitness due to selection in evolutionary population dynamics. The ability of a community to change in a directional way and that the same change favors its expansion in space (“reproduce”), can be viewed as a Lamarkian feature [11].

The behavior of the meta-community changes qualitatively between early and late times. At early times, it develops by a complex combination of internal changes and cross-community influences, different facets of which can been described in multiple ways. A process in which a new state forms by combining the previous community state and that of its neighbors, rather than a perfect copy of a single state, has parallels with sexual reproduction [41, 48]. At late times, however, the dominant mode of change in the system starts with a jump to a new state in one community, which then replicates faithfully across the entire system, see Fig. 6(A,C) and additional examples in the Supporting Information, Appendix B. Such behavior is analogous to a mutation and replication process, with a mutation that is, a priori, directional rather than neutral. This “mutation-replication” behavior is an outcome of the dynamics, rather than a model assumption.

The growth of stability described in Sec. III, in the sense that communities spend ever longer times at a single state or near it, means that changes occur over many different time scales, from time scales of an individual species response to many orders of magnitude above that. In meta-communities, the dynamics might speed up significantly with the number of communities, as states which might take over are explored in parallel in the different communities. When studying systems over long time scales, additional processes such as environmental fluctuations or evolution might become important and should be taken into account. Introducing evolution may raise interesting questions regarding multi-level selection [54], once three levels of organization-genes, organisms and communities-play an active role.

### Theoretical implications and comparison with other works

Comparing with previous work, theoretical mechanisms for directionality often require strong hierarchies or fine-tuned trade-offs between species, that might not emerge in a high-dimensional space of traits and interactions [28, 36]. Some models are directional by construction, for example certain resource competition models [33, 34] but not others [26]. In MacArthur’s resource-competition model [33, 34] and related models [49], there is a single species composition that is stable to invasions. Here, in contrast, there are many uninvadable states, and the jumps between them are responsible for the emergence of long time-scales in the dynamics. Many other, different quantities have been proposed to grow during succession (maximum principles); their origins and regimes of applicability continue to be debated [18].

Some models do have many alternative equilibria, and have a function that grows in time; however, they require symmetric interactions, are sensitive to chaos when the symmetry breaks [43], as is shown in Fig. 2.

We remark that the multiple equilibria discussed here are different from regime shifts [3, 4, 6, 29, 45, 51], in which large qualitative changes occur in the ecosystem. Here, in contrast, there may be very many closely-related alternative stable states, which differ in the details of the species composition, but may have similar system-wide properties such as species richness or functioning.

The work raises a number of interesting directions for future theoretical investigation. Firstly, while we have proposed some intuitive explanations for the observed phenomena, a deeper theoretical understanding is called for, with the goal of understanding the entire dynamics within a single coherent picture. Why does the maturity *F* increase, and what is its functional form? We show that the phenomena described here are robust under certain changes to the model (see Supporting Information, Appendix A), and also that there are limits to the region of phase space where a qualitatively different behavior takes over, for example persistent abundance fluctuations (see Sec. II), or when noise is strong and washes out the directionality (see Supporting Information, Appendix A). It would be interesting to understand the location and characteristics of the boundaries between these different regions (’phase transitions’), as discussed in Sec. VII. Another aspect of the model is the sparsity of interactions, where each species significantly interacts with only a few other species, whose role remains to be understood.

It would be interesting to further explore other situations where these phenomena can be found. For example, mutually exclusive pairs might be replaced by groups of species that compete in a more structured interaction pattern, without any specific pair being mutually exclusive. In the present work, the interactions are predominantly competitive. Can similar models be constructed for strong (obligate) mutualism, an important driver of close associations between species [11, 12]?

Another interesting question is whether a quantitative description be constructed at the level of the *F*-values of states alone, without reference to additional details? For example, how well does the *F*-value of a given state predict its ability to expand, and why?

### Conclusion

Our theoretical investigation paints an intermediate picture of succession, between the extremes of developmental stages [14] and chance combinations of species [24]. While strictly ecological, this picture can be better understood by borrowing concepts from evolutionary dynamics. We summarized above multiple fingerprints that could hopefully allow to identify this process in natural communities. We have shown that observing many species associations and stochastic transitions does not necessarily stand against directionality and community-level The ability to measure and control such interactions in the laboratory [1, 37, 56] gives hope that the theoretical predictions can be directly tested.

## IX. MATERIALS AND METHODS

Here the basic variant of the model, used above, is defined. For convenient reference, all parameters used in all simulations are concentrated here, including those already quoted above.

The interaction matrix *α* in the model used in Sec. II,III,IV,VI, is constructed by first sampling a network of non-zero interactions. It is taken to be a random regular graph, i.e. where each species interacts with *C* other species, generated using the algorithm in [46]. For a pair of species *i, j* that interact, both the interactions *α_ij_, α_ji_* are non-zero and sampled independently from a Gaussian distribution.

The parameters are as follows: In Fig. 2(A), *C* = 20, std (*α_ij_*) = 0.4, *S* = 400, and all *r_i_* = 1. In Fig. 3,4 and 6, *C* = 3, mean (*α_ij_*) = 1.667, std (*α_ij_*) = 0.75, *S* = 24. The growth rates *r_i_* were sampled independently for each *i*, from a uniform distribution on [0.5, 2]. Everywhere, *λ_i_* = 10^−5^ for all *i*. The noise amplitude is *ω_i_* = 0.173 for all *i* in Fig. 3,4. In Fig. 6, *ω_i_* = 0.1 and *D* = 0.1. Initial conditions for *N_i_* (0) are uniform on [0, 1].

The *S* = 2 models in Fig. 2, have *α*_12_ = 4, *α*_21_ = 2 in and *α*_12_ = 0.6, *α*_21_ = 0.4 in (C). The 3-species model shown in Fig. 5 is defined by *α*_12,21,13,31_ = 1.5,*α*_23,32_ = 0, *ω_i_* = 0.173, *r_i_* = 1 and *D* = 0.03.

Zero-noise dynamics shown in Fig. 2 are simulated with migration rate *λ* = 10^−15^, using a Runge-Kutta ordinary differential equation solver, with relative and absolute error tolerance of 10^−13^.

Noisy dynamics are integrated by a standard stochastic differential equation algorithm. The abundances *N_i_* (*t*) are advanced by

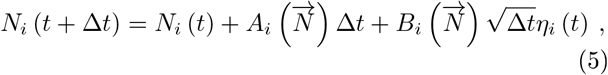

where 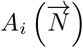 is the deterministic term in the right-hand side of Eq. (1,4), and 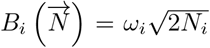 is the prefactor of the white noise, and *η_i_* (*t*) is a Gaussian random variable with unit variance and zero mean, sampled independently for each species and time step. We take Δ*t* = 0.01. If some *N_i_* (*t* + Δ*t*) goes below 10^−10^, it is set to 10^−10^ (with *λ* = 10^−5^ used in all stochastic simulations, this doesn’t happen often). A different regularization, 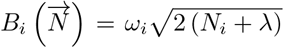, has also been tested and gives very similar results.

Correlations between two abundances at two times, 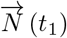, 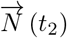 in one community are calculated by first recording

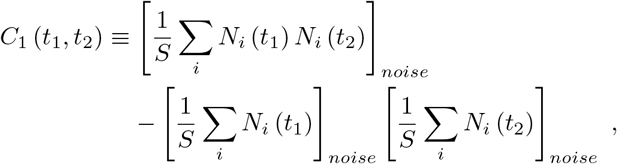

where [..]_*noise*_ denotes the average over noise realizations with the same initial conditions and *α_ij_, r_i_*, where 10 repetitions with different noise realizations are used. Averaging *C*_1_ (*t, t*′) over model samples (*α_ij_, r_i_*) and initial conditions gives *C_u.n._* (*t, t*′). Finally, the correlation coefficient shown in the main text is given by: 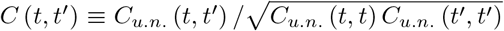. In metacommunities, the correlations for different communities are computed similarly, only with 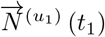, 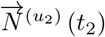 where *u*_1_, *u*_2_ represent different communities, and the sum over *i* is replaced by a sum over *i* and all *u*_1_ ≠ *u*_2_.

## Acknowledgments

It is a pleasure to thank J.F. Arnoldi, M. Barbier, J. Friedman and M. Loreau for helpful discussions and critical reading of the manuscript. Support by the Israel Science Foundation (ISF) Grant no. 773/18 is gratefully acknowledged.

## Appendix A: Variants of the basic model

The main properties discussed in the main text are insensitive to many of the model’s details. We now discuss a number of variants of the basic model. For all of them we show ⟨*F* (*t*)⟩, the equivalent of the smooth line in Fig. 3(B). For five variants of the model, we show that ⟨*F* (*t*)⟩ continues to increase over long times, roughly as ⟨*F* (*t*)⟩ ∝ ln *t*, as in the basic variant of the model used in the main text.

At the same time there are phase boundaries that, when crossed, may change the behavior significantly, as in the transition from directionality to chaos upon changing *α_ij_* discussed in Sec II, see Fig. 2. Below we also give another example of such a transition, when the noise is increased beyond some value.

The basic model variant used in the main text has a number of mutually-exclusive pairs of interactions per species. Other interaction-types may be present too. Already in the basic model (Figs. 3,4 and 6), 66% of the the non-zero interaction pairs are mutually-exclusive (both *α_ij_, α_ji_* > 1), 30% have just one *α_ij_* > 1, and the rest may coexist in pairs (*α_ij_, α_ji_* < 1). In a first variant, we look at graphs where 15% of the interactions are directional, such that one of the interactions is non-zero (*α_ij_* > 0 with *α_ji_* = 0). This is done by creating a random regular graph (undirected) with *C* = 3 links per species and another directed graph in which each species *i* has 3 *j*-values for which *α_ij_* is non-zero (*j* = *i* allowed). Then, edges from the former network are taken with probability 0.85 and from the latter with probability 0.15 (this also reduces the fraction of non-zero mutually-exclusive pairs to 57%). Running simulations with these interactions shows a similar behavior of *F* (*t*), see Fig. A(A), “additional directed links”. (Other model parameters: *S* = 24, mean (*α_ij_*) = 1.667, std (*α_ij_*) = 0.58, *ω_i_* = 0.173, *r_i_* uniform on [0.5, 2]).

In another variant, interactions are taken to have a non-linear form,

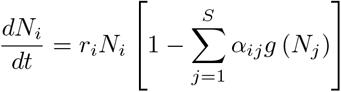

with a saturating Holling type-II function for the interactions, 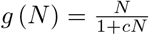. This requires a different formula for the growing function *F*. Indeed, in the case where the matrix *α* is symmetric, the Lyapunov function generalizes straight forwardly to [40]

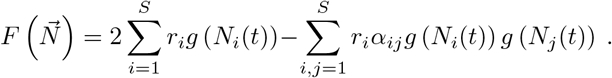

**Figure A.**
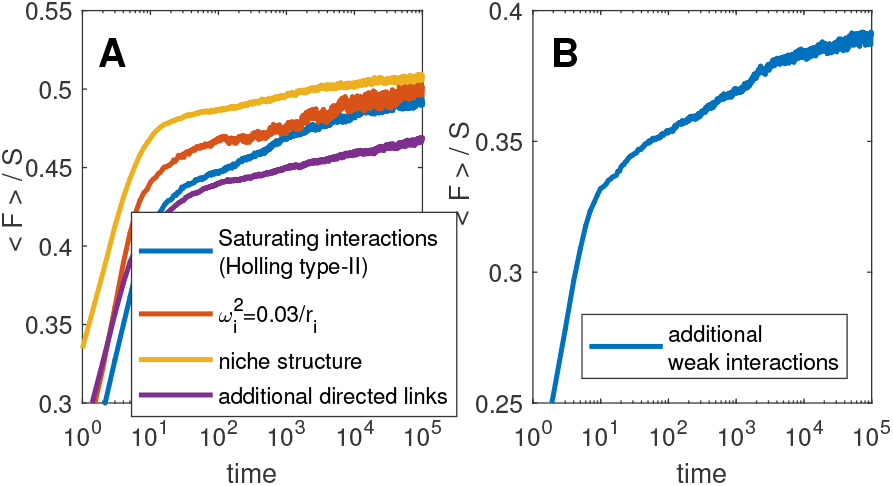
⟨*F* (*t*)⟩ for different model variants. Full definitions and parameters values are given in the text.

The behavior of ⟨*F* (*t*)⟩ with this function also increases approximately logarithmically, see Fig. A(A), “saturating interactions”. (Model parameters: *S* = 24, *C* = 3, mean (*α_ij_*) = 1.667, std (*α_ij_*) = 0.58, *ω_i_* = 0.173, and *c* = 1, *r_i_* uniform on [0.5, 2]).

Another variant regards the network of non-zero interactions. In the basic model it is sampled as a random regular graph, with 3 links per species. Here we sample it from an underlying niche structure. The different species are embedded in a two-dimensional niche space, in which each species’ location is a point sampled uniformly in the square (0, 1) × (0, 1). Two different species are assigned a non-zero interaction if the distance between them is smaller than *d*_0_. We choose *d*_0_ = 0.23, so that the *average* number of non-zero pairs is about 3, but this number fluctuates significantly between species. Moreover, this construction gives the interaction network additional structure beyond a random regular graph. For example, the network has some degree of modularity [?]: if species *i, j* are close in niche space so as to interact, as are species *j, k*, then it is more likely that species *i, k* are close enough to interact. With this structure, the logarithmic growth is still observed, see Fig. A(A), “niche structure”.

Another change involves having noise amplitudes *ω_i_* that are dependent on *i*. If these are random numbers that are not correlated with any other system parameters, little changes in the behavior. Interestingly, the logarithmic growth survives even if *ω_i_* has some relation to *r_i_*, here tested for 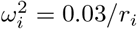, see Fig. A(A) (Parameters: *S* = 24, *C* = 3, mean (*α_ij_*) = 1.667, std (*α_ij_*) = 0.75, *r_i_* uniform on [0.5; 2]).

Finally, we demonstrate the effect of additional weaker interactions in all pairs. The model takes the interactions in the basic model *S* = 24, *C* = 3, mean (*α_ij_*) = 1.667, std (*α_ij_*) = 0.75, and adds to them non-zero inter-actions to all pairs *i* ≠ *j*, sampled independently from *P* (*α_ij_*) = *e*^−*α_ij_*/0.04^, so that the mean of the additional terms is 0.04. Note that while individually these numbers are small compared to the nominal interactions, they appear for *all* interactions pairs, rather than just for *C* = 3 links per species. Here too ⟨*F*⟩ grows over long time scales, see Fig. A(B).

**Figure B.**
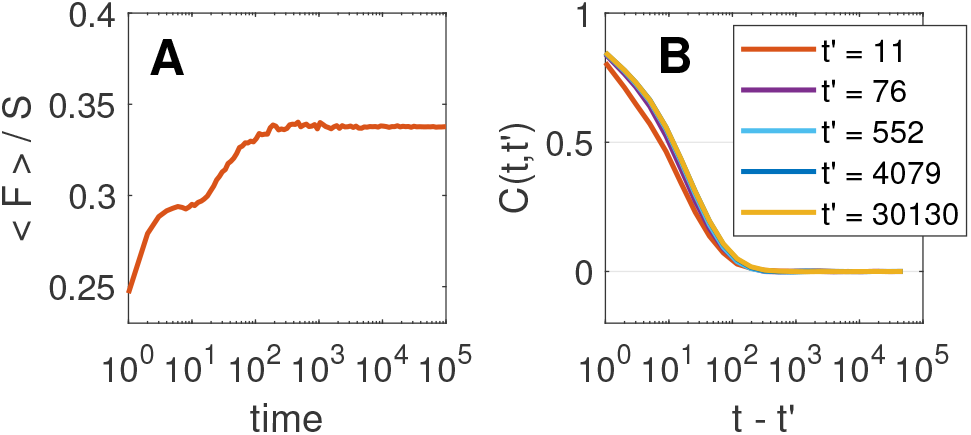
The equivalent of Fig. 3(B,C) in the same model, only with larger noise (*ω_i_* = 0.4). (A) The function ⟨*F* (*t*)⟩ increases at first but saturates and does not increase at long times. (B) The correlation *C* (*t, t*′) decays after shorter times (*t* − *t*′ ~ 10^2^) and for late times depends only on the time difference *t* − *t*′, as shown by same behavior for *t*′ = 4079, 30130. These results mean that here there is no long-time change in the dynamical behavior, that could come from transitions towards states with ever higher stability.

For comparison, it is instructive to see an example that shows a different behavior. Here we look at a system exactly like that in Fig. 3,4 and 6, only with higher noise, *ω_i_* = 0.4 (instead of 0.173), see Fig. B. The function ⟨*F* (*t*)⟩ grows but saturates at times *t* ~ 100, and does not show a subsequent logarithmic growth (in contrast to Fig. 3(B)). Together with that, the correlation *C* (*t, t*′) from time *t*′ to *t* decays more quickly (compare with Fig. 3(C)). Crucially, at later times *t*′ it depends only on the time difference *t* − *t*′, i.e. *C* (*t, t*′) = *f* (*t* − *t*′), as seen in the almost identical dependence on *t* − *t* of the correlations starting at *t*′ = 4079, 30130.

While growth of *F* (*t*) implies reaching a fixed point, the converse is not generally true. For example, a fixed point is reached in Fig. 2(B,C) but *F* (*t*) decreases on the way, and similarly for high *S* dynamics discussed in [13], in which a fixed-point is reached.

## Appendix B: Higher diversity and correlations between communities

In this section the results at two different diversities *S* are compared. It also shows the cross-correlation between communities, not shown in the main text.

As the number of species *S* is increased, the mean ⟨*F* (*t*)⟩ */S* converges to a function, quite close to the the one for *S* = 24, see the comparison with *S* = 100 in Fig. C(A). Individual runs with higher *S* follow the mean ⟨*F* (*t*)⟩ more closely, and have more transitions between states, see example in Fig. C(A). The two-time correlation *C* (*t, t*′) converges to a well-defined limit as *S* grows, so the increase in the number of transitions does not induce a faster decorrelation; Again, the function for *S* = 24 is quite close to the asymptotic function. For any *S* (as for *S* = 24), the rate of transitions between states strongly decreases with time, and so even at higher diversity there is a time after which the system spends extended periods of time in a given state, punctuated by transitions to new states.

**Figure C.**
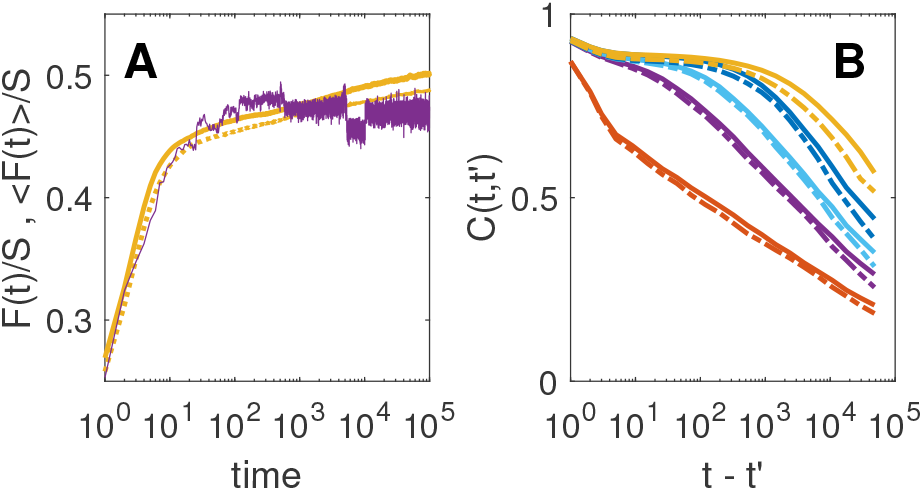
(A) ⟨*F* (*t*)⟩ for *S* = 100 (thick solid line), compared with the same for *S* = 24 (dotted line), as in Fig. 3(B). A single example run for *S* = 100 is shown. (B) The two-time correlation *C* (*t, t*′) for *S* = 100 (solid lines) and *S* = 24 (dotted lines). Line of different colors show different *t*′ values, as in Fig. 3(C).

**Figure D.**
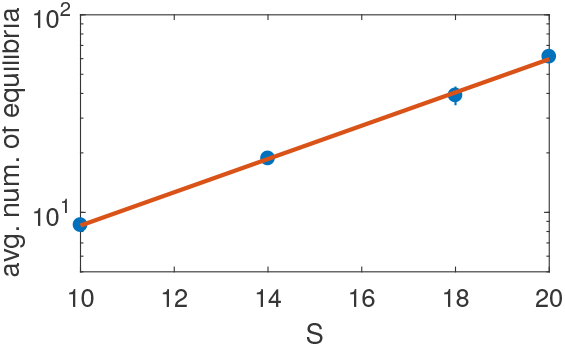
Average number of stable equilibria as a function of *S*. Solid line: fit to exponential, *e*^0.194*·S*+0.209^. Parameters as in Fig.3,4 and 6 (*C* = 3, mean (*α_ij_*) = 1.667, std (*α_ij_*) = 0.75).

The number of equilibria increase with *S*, see Fig. D. They are counted by exhaustively checking all subsets of species for equilibria. The dependence fits very well to an exponential between *S* = 10 to 20 (a dependence that was shown for a related, more stylized model [21]). This predicts that at *S* = 24 there will be on average about 130 states, and for *S* = 100 about 3 10^8^ states. While there are many equilibria, these only represent a small fraction of all possible states (for example, the number of subsets of species for *S* = 100 is 2^100^ ≃ 10^30^). Thus, most subsets of species are not valid equilibria, in contrast with the idea of completely random species associations.

Cross correlation between neighboring communities in a meta-community are shown in Fig. E. Parameters are the same as for Fig. 6, with *D* = 0.1. The correlation between nearby communities is less than one even if the communities are fluctuating in the same state, because they experience different demographic noise in each community. Results for *S* = 24, 100 are shown, and again found to be very similar.

It is interesting to note that the spread of states through the meta-community causes any given community to experience more changes of state, as the new states can be created in any community before spreading. To test this, we looked the state of one community within the meta-community, at *S* = 24, at times 30, 100, 300, 10^3^, 3 · 10^3^, 10^4^, in repeated runs with parameters as in Fig. 6(A). The number of changes of states between subsequent times was on average larger by 43%_(±3%)_ than for a single isolated community.

**Figure E.**
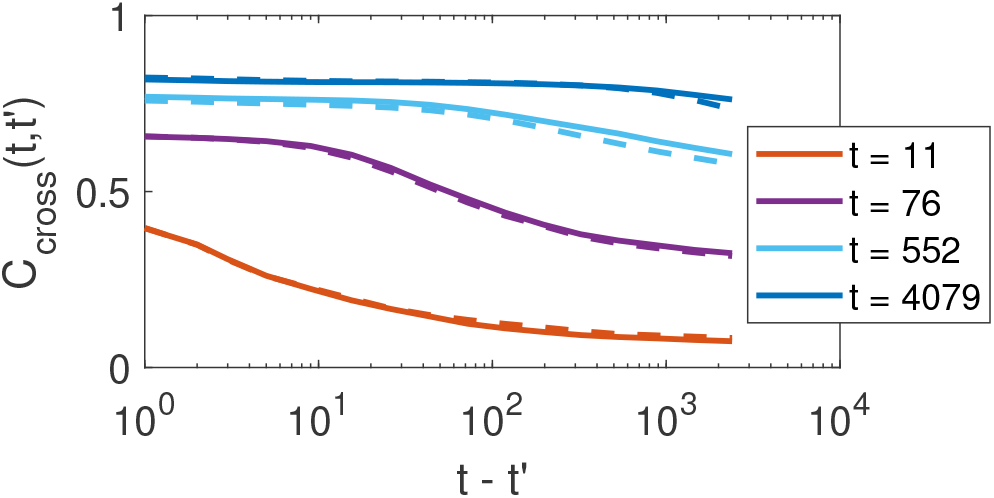
Cross-correlation between neighboring communities in a chain of *M* = 10 communities as in Fig. 6. Comparison between *S* = 100 (solid lines) and *S* = 24 (dashed lines).

*Additional run samples*-Fig. F shows 4 runs sampled as in Fig. 6(A), except for *S* = 100 here. The four runs are an unbiased sample-without any additional selection by the author. Fig. G shows a close up on two areas in Fig. F, showing sweeps in which a new state takes over the system. Fig. F(II) shows two consecutive sweeps; in the later a state with somewhat lower *F* takes over.

**Figure F.**
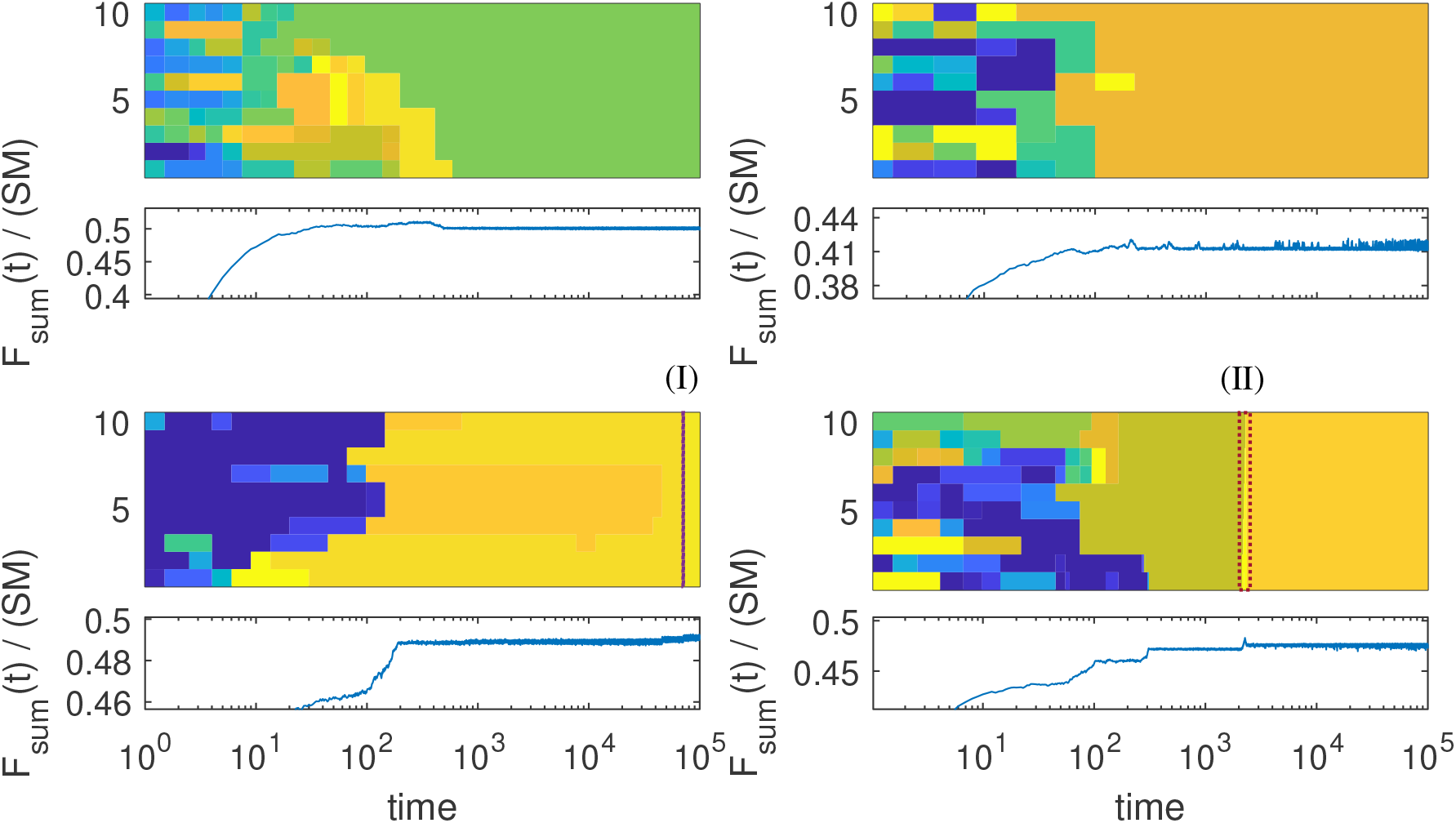
Additional run samples, with the same model parameters as Fig. 6(A-D) except for *S* = 100. For each run the map of states is shown as in Fig. 6(A), and below each *F* (*t*) / (*SM*) is plotted as in Fig. 6(B). In order to show difference at late times, the colors do not differentiate all the early states with low *F*-values.

**Figure G.**
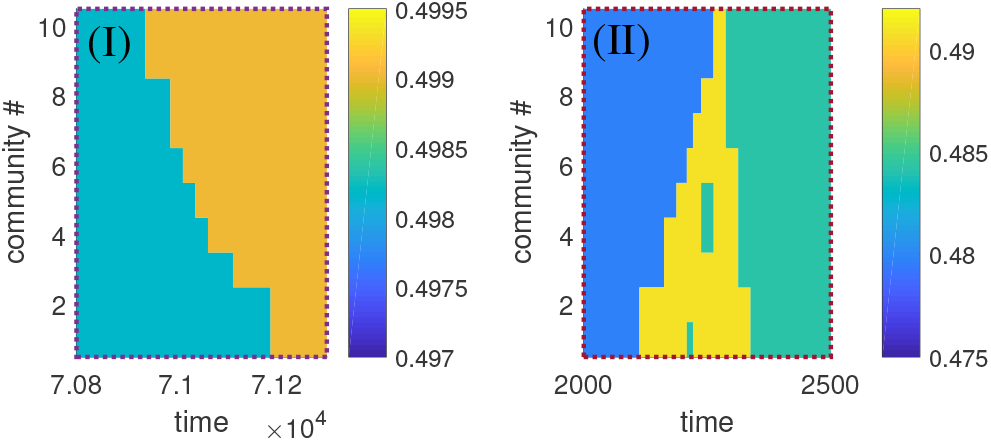
Close-up on two areas in Fig. F. For clarity, colors have been changed from Fig. F.

## Footnotes

1 To admit a Lyapunov function, the matrix *A* can be of a somewhat more general form than symmetric [40]. The model described below does not satisfy this form.

2 This behavior depends on the parameter 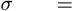 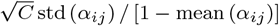 [10, 13], which diverges at mean (*α_ij_*) = 1. A transition to stable equilibria has also been reported in simulations of [27].

3 In some runs with ⟨*α_ij_*⟩ = 1.5 there may be a small decrease in *F*, with minimal *dF/dt* of order −10^−5^. This small effect is suppressed when ⟨*α_ij_*⟩ is higher (say, ⟨*α_ij_*⟩ = 2) and λ smaller.

4 This is in contrast with simple models of multi-stability which only contain two (or a handful) of alternative states [44], where *C* (*t, t*′) would depend only on the time difference *t* − *t*′, decaying after the time typical to the jump between these states. The behavior observed in the present model is characteristic of dynamics in complex high-dimensional landscapes [2, 9]. It is also distinct from the slow-down of recovery *within* a state from typical noise-induced fluctuations, when an equilibrium is almost unstable [45]. In contrast, here the initial decay of *C* (*t, t*′) (at *t − t*′ ≲ 10) corresponding to relaxation within a state, is similar for all *t*′ ≥ 76, see Fig. 3(C).

